# Ultra-accurate Genome Sequencing and Haplotyping of Single Human Cells

**DOI:** 10.1101/135384

**Authors:** Wai Keung Chu, Peter Edge, Ho Suk Lee, Vikas Bansal, Vineet Bafna, Xiaohua Huang, Kun Zhang

**Author notes:** To whom correspondence should be addressed.; or Data deposition: Sequence data are available at NCBI SRA (SUB2632622) Code is available at https://github.com/pjedge/SISSOR.

## Abstract

Accurate detection of variants and long-range haplotypes in genomes of single human cells remains very challenging. Common approaches require extensive *in vitro* amplification of genomes of individual cells using DNA polymerases and high-throughput short-read DNA sequencing. These approaches have two notable drawbacks. First, polymerase replication errors could generate tens of thousands of false positive calls per genome. Second, relatively short sequence reads contain little to no haplotype information. Here we report a method, which is dubbed SISSOR (Single-Stranded Sequencing using micrOfluidic Reactors), for accurate single-cell genome sequencing and haplotyping. A microfluidic processor is used to separate the Watson and Crick strands of the double-stranded chromosomal DNA in a single cell and to randomly partition megabase-size DNA strands into multiple nanoliter compartments for amplification and construction of barcoded libraries for sequencing. The separation and partitioning of large single-stranded DNA fragments of the homologous chromosome pairs allows for the independent sequencing of each of the complementary and homologous strands. This enables the assembly of long haplotypes and reduction of sequence errors by using the redundant sequence information and haplotype-based error removal. We demonstrated the ability to sequence single-cell genomes with error rates as low as 10^−8^ and average 500kb long DNA fragments that can be assembled into haplotype contigs with N50 greater than 7Mb. The performance could be further improved with more uniform amplification and more accurate sequence alignment. The ability to obtain accurate genome sequences and haplotype information from single cells will enable applications of genome sequencing for diverse clinical needs.

## Significance

Accurate sequencing and haplotyping of diploid genomes of single cells are intrinsically difficult due to the small amount of starting materials and limited read lengths of current DNA sequencing methods. In SISSOR, we aim to improve sequencing accuracy and haplotype assembly by taking advantage of the redundant complementary sequence information in the double-stranded DNA, and by partitioning megabase-size single-stranded DNA fragments from the homologous chromosome pairs into multiple compartments for amplification by MDA (multiple displacement amplification), and subsequent sequencing using short-read DNA sequencing platforms. We report the demonstration of this concept using sequence data from three single human cells. Our approach can simultaneously provide higher accuracy and longer haplotypes than existing approaches.

The ability to accurately identify variants in both coding regions and functional regions is essential to clinical genomics. Single nucleotide polymorphisms (SNP) are the most common type of genetic variations. SNPs are estimated to appear in about every 100-300 bases and account for 90% of all human sequence variations (1, 2). The abundance of SNPs also provides the major source of heterogeneity for linking variants in a haplotype, where the combination of alleles occurs at multiple loci along a single chromosome. The ability to confidently identify *de novo* somatic mutations on top of myriads of SNPs in single mammalian cells is challenging. Current single-cell genome sequencing approaches can manage to call millions of germline variants but also generate tens of thousands of false positive calls that greatly outnumber the somatic mutations per genome. Besides the detection of *de novo* mutations, accurate long-range haplotyping is also useful for many clinical applications. For example, the human leukocyte antigen (HLA) haplotype spans a ∼5 Mb region in human chromosome 6. The HLA genes, which encode cell surface proteins and regulate the immune system in humans, tend to inherit as a cluster within a single haplotype. Accurate HLA haplotyping allows for better donor-patient matching for organ transplants.

Unlike targeted sequencing such as exome sequencing, whole genome sequencing (WGS) allows for the detection of all known and unknown variants. WGS using current DNA sequencing technologies requires input DNA equivalent to tens to thousands of cells. Therefore, whole genome sequencing of single cells invariably requires amplification of genomic DNA. Unfortunately, many errors are introduced in the amplification process, with error rates ranging from 1.2 × 10^−5^ in conventional MDA to as high as 2.1 × 10^−4^ in MALBAC (multiple annealing and looping-based amplification cycles) (3). In addition, current DNA sequencing technologies have substantial error rates and limited read lengths (4–6). Long-range haplotype information is very difficult or impossible to be obtained from the short sequences provided by highly parallel short-read sequencing platforms (7). The current approaches to achieve a haplotype greater than 1 Mb rely on methods to haplotype single DNA molecules. These strategies require extensive preparation in cloning (8), isolation of metaphase chromosomes (9), segregation of complementary strands in dividing daughter cells (10), or parallel partitioning of genomic DNA in water-in-oil microdroplets (11). These limitations make accurate single-cell whole genome sequencing and long-range haplotyping very challenging.

Here we introduce a new method called SISSOR, in which double-stranded chromosomal DNA molecules from single cells are separated and megabase-size fragments are stochastically partitioned into multiple nanoliter compartments for enzymatic amplification in a microfluidic device. The random partitioning allows for independent amplification and sequencing of the homologous chromosomal DNA fragments for long-range haplotype assembly and error correction by comparing the phased complementary strands. As a proof of concept, we amplified and sequenced three single cells from a human fibroblast cell line (PGP1f) whose genome has been sequenced extensively using other approaches.

## Results and Discussion

### Partitioning and Amplification of Separated Single-stranded Chromosomal DNA Fragments

We implemented the SISSOR concept using an integrated microfluidic processor. The device and the overall procedure are illustrated in Figure 1 (more detail in Figure S1 in SI Appendix). The microfluidic device consists of four modules: single-cell capture, cell lysis and strand separation, partitioning, and amplification modules. A single cell is captured from a cell suspension. The cell is lysed and double-stranded chromosomal DNA molecules are separated using an alkaline solution. The separated DNA fragments are randomly distributed using a rotary pump and partitioned into 24 identical chambers. Each partition is then neutralized and pushed down into the amplification module and amplified by MDA. Amplified products in each chamber are retrieved, converted into barcoded sequencing libraries and sequenced using Illumina short-read sequencing-by-synthesis.

**Fig. 1.**
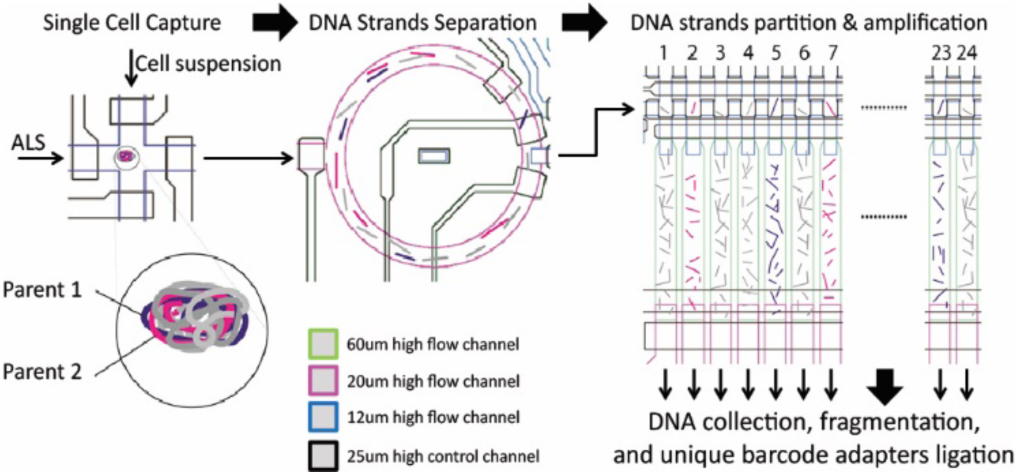
An overview of the experimental process of SISSOR technology. A single cell in suspension was identified by imaging and captured. The cell was lysed and chromosomal DNA molecules were separated into single-stranded form using an alkaline lysis solution (ALS). The single-stranded DNA molecules were randomly distributed and partitioned in 24 chambers. Each partition was pushed into an air-filled MDA chamber using a neutralization buffer, followed by an MDA reaction solution. MDA reaction was carried out by heating the entire device at 30 °C overnight. The amplified product in each individual chamber was collected out of the device and processed into barcoded sequencing library.

We experimented with various procedures and designs of the processor to optimize SISSOR. We found that cell lysis, denaturation of the long double-stranded chromosomal DNA molecules, and the distribution of the ssDNA fragments were more effective with a higher KOH concentration (up to 400 mM) and pumping speeds of the rotary mixer. To prevent DNA damage at high pH, the process was limited to 10 minutes at 20 °C. We also found that MDA chambers with a lower surface to volume ratio and precoating of the polydimenthylsiloxane (PDMS) surface with bovine serum albumin improved the amplification, perhaps by reducing the inhibition of the DNA polymerase and the non-specific binding of molecules. MDA chambers with a volume of ∼20 nl provided sufficient amplification with less bias by limiting the available nucleotides (dNTPs).

#### Genome Coverage

We amplified the genomes of three single cells from human PGP1 fibroblast cell line using the optimized devices and procedures. The amplified genomic DNA collected from each of the individual 24 chambers was converted into barcoded sequencing libraries. The barcoded libraries from each cell were combined and sequenced using standard 100 bp paired-end Illumina sequencing. Sequencing reads from the individual chambers were identified using the barcodes, and mapped to the reference human genome hs37d5 (GRCh37/b37 + decoy sequences) using the default setting of BWA-MEM with Burrows-Wheeler Aligner (BWA) (SI Appendix, Fig. S2A). Deep sequencing data from the three single cells yielded a combined 558 gigabases (Gb), with 92.6%-98.8% mappable reads. We obtained an average of 65-fold sequencing depth and 63.8%±9.8% genome coverage per cell. The combined sequence reads from three cells cover 94.9% of the entire genome (SI Appendix, Table S1). The high mapping rate was perhaps the result of the high fidelity Phi-29 polymerase used in MDA and reduced contamination in small reaction volumes of the microfluidic devices. BWA-MEM also identified and removed chimeric reads commonly introduced by MDA. Since the combined genome coverage from three cells is ∼30% higher than any individual cell, we suspect that some fragments were lost during strand separation and partitioning within the device and did not reach the amplification chambers.

#### Determination of SISSOR Fragments

Megabase-sized DNA fragments were visualized by mapping all sequencing reads to the reference genome sequence (SI Appendix, Fig. S3). The individual DNA fragments appeared as dense blocks with overlapping unique reads. Genome regions that have dense mapped coverage with reads from two or more chambers indicate that the four complementary strands of the homologous chromosome pairs were separated, partitioned and amplified in different chambers (Fig. 2). The reads from each chamber that are mapped to sporadic regions of the genome with very low density coverage are very likely the consequence of mis-alignment. Thus, they were removed in the segmentation of SISSOR fragments using hidden Markov model (SI Appendix, Supplementary methods).

**Fig. 2.**
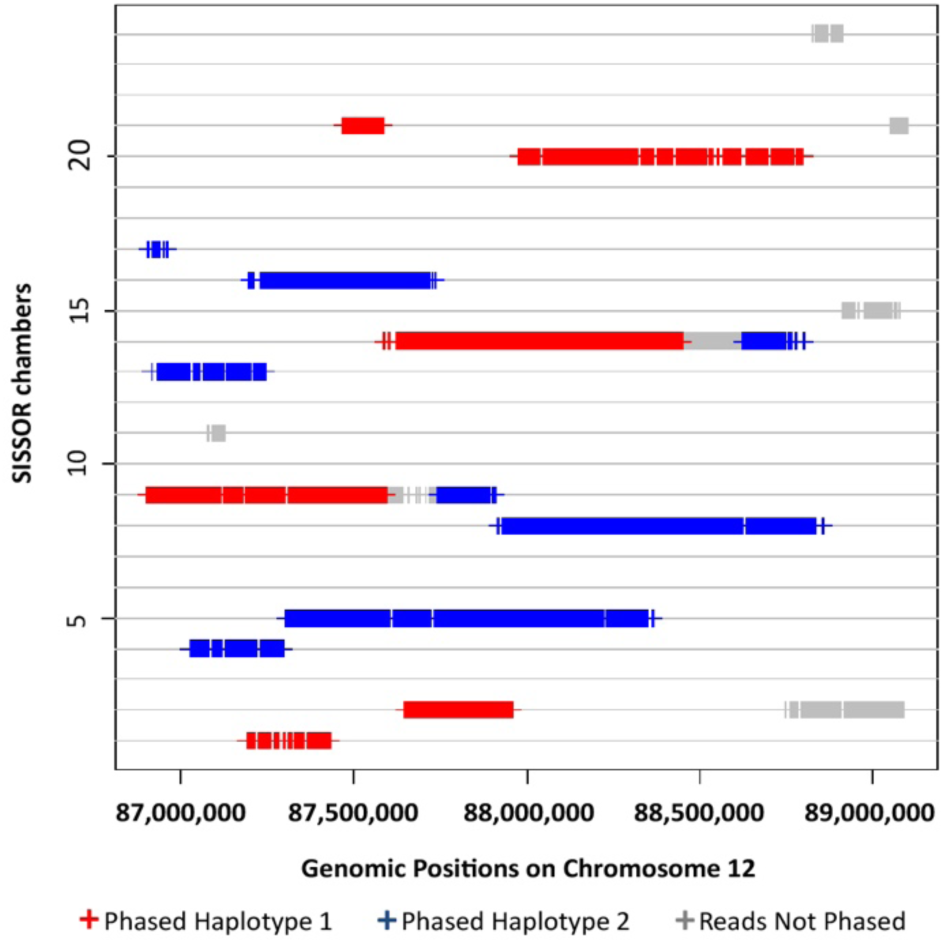
Haplotyping of single-stranded DNA fragments using sequencing reads from single-cell genome amplified using a SISSOR device with 24 chambers. Large sub-haploid SISSOR fragments were first computed per chamber and then phased into haplotype 1 and haplotype 2 with HapCUT2 (12). SISSOR fragments could be visualized by mapping the sequencing reads to a reference genome. Some fragments were not phased due to either the lack of heterozygous SNPs or the presence of mixed sequences from two or more strands.

We determined the boundaries of the large SISSOR fragments by joining the aligned sequencing reads using an HMM, based on read depth and proximity in the localized genomic regions. We counted the number of reads in each good bin defined by the 50K variable bin method (13) and calculated the average number of reads if these reads were randomly distributed in all 50K bins (SI Appendix, Figs. S2B and S4). The resulting fragment boundaries as determined by the start and end positions of continuous bins in HMM were highly consistent in the range of 1x-5x average reads per bin. These boundaries closely resembled the sub-haploid DNA fragments because of the high ratio of reads per bin concentrated in a small genomic region rather than distributed randomly in the entire genome. About 11.8% of mapped locations were removed in HMM by choosing 5x average reads per bin (SI Appendix, Table S2). The N50 fragment size exceeded 1 Mb with the largest contig at 9 Mb (SI Appendix, Table S3). The mean DNA fragment length prior to haplotype assembly is ∼500 kb, which is 5-10 fold longer than what has been achieved using dilution methods. We estimated that each single-stranded DNA fragment was amplified about 60,000 times.

#### Variant Calling

The unique design of SISSOR enables more accurate single-cell variant calls in two ways: first, variants that are observed in multiple chambers can be called more confidently than variants observed only once (multiple chamber allele occurrence). Second, variant calls that match between strands from the same haplotype are of especially high confidence (same-haplotype strand matching).

To leverage multiple chamber allele occurrence, we developed a novel variant calling algorithm. The algorithm closely models the SISSOR workflow to make consensus allele calls for every genomic position using sequence information from all chambers (SI Appendix, Figs. S2C, S2D and Supplementary methods). Briefly, the algorithm models all the possibilities that single DNA strands from a diploid genome could be distributed to and amplified in the chambers, and accounts for possible error sources such as MDA. The algorithm assigns higher confidence to alleles that are observed multiple times and fit well into the diploid model. We gauged the accuracy of our algorithm at different confidence thresholds by comparing to a reference genome sequence for PGP1 (SI Appendix, Table S4, and Supplementary methods). At the most lenient threshold, 1.7 million SNVs were called with a false positive rate of 5×10^−5^. At a moderate threshold, 613,669 SNVs were called with a false positive rate of 1x10^−6^. At the strictest threshold, 177,096 SNVs were called with a false positive rate of 1×10^−7^.

Even greater accuracy can be achieved by leveraging same-haplotype strand matching, an approach which requires separating fragments into different haplotypes. To perform haplotype assembly, we extended our variant calling model to call the most likely allele in every chamber (at a lenient threshold) and generate sub-haploid fragment sequences (SI Appendix, Supplementary methods). In the following sections, we describe haplotype assembly and validation of variant calls by same-haplotype strand matching to achieve maximum accuracy using the SISSOR technology.

#### Whole Genome Haplotyping

Haplotype assemblies were constructed by phasing heterozygous SNPs in sub-haploid SISSOR fragments. A list of heterozygous SNPs, obtained from 60x coverage Illumina WGS data of PGP1 fibroblast cells (under ENCODE project “ENCSR674PQI”), was used to phase the 1.2 million SNPs in SISSOR fragments. We applied these SNPs to a haplotyping algorithm, HapCUT2 (12), and compared the assembly to the PGP1 haplotype created using sub-haploid pools of BAC clones (8). Two types of errors may occur in an assembled haplotype. First, a switch error was defined as two or more SNPs in a row flipped. Second, a mismatch error was defined as a heterozygous SNP whose phase was incorrectly inferred. If a higher switch and mismatch error rate (1.6%) could be tolerated in an application, a large N50 haplotype length (> 15 Mb) was directly produced by HMM-derived SISSOR fragments. We anticipate that genome resolution can be augmented by mapping high-quality short sequencing reads to the long haplotype scaffold. Similarly, long-range chromosome-length haplotype scaffolds have been created with the Strand-seq approach, which required BrdU incorporation in dividing cells (10). Combining the heterozygous variants in short WGS reads (∼250 bp) to long haplotypes was shown to improve the phased coverage.

We further processed and refined the raw SISSOR fragments to address the case where two overlapping homologous DNA fragments may appear in the same chamber (SI Appendix, Fig. S2D). Long SISSOR fragments were split where the phase of two SNPs in a row are flipped with respect to fragments from other chambers. We removed the fragments with clusters of low quality variant calls, and then reassembled these processed fragments with HapCUT2. Splitting longer fragments with detecTable Switch errors and poor variant calls from mixed homologous reads at the unique genomic position reduced the overall haplotyping errors. Four-strand coverage of processed fragments reduced more than 17% of the original size but the phasable whole SISSOR fragments increased from 70-80% to about 93% in all three cells.

Although the lengths of processed SISSOR fragments were reduced, HapCUT2 assembly of overlapping fragments still creates long haplotype contig with an N50 ∼7 Mb and with >90% span of the human genome (Table 1). In comparison to BAC haplotype, which has an N50 ∼2.6 Mb for the PGP1 cell line, the 1.2 million heterozygous SNPs called and phased in our SISSOR libraries have a concordance rate of 99.3% in our assembled haplotype blocks. The achieved haplotype concordance in SISSOR is comparable to the 99.1% and 99.4% accuracy rates obtained using fosmid clone and PacBio 44x coverage SMRT data, respectively, for the NA12878 genome using HapCUT2 (12).

**Table 1.**
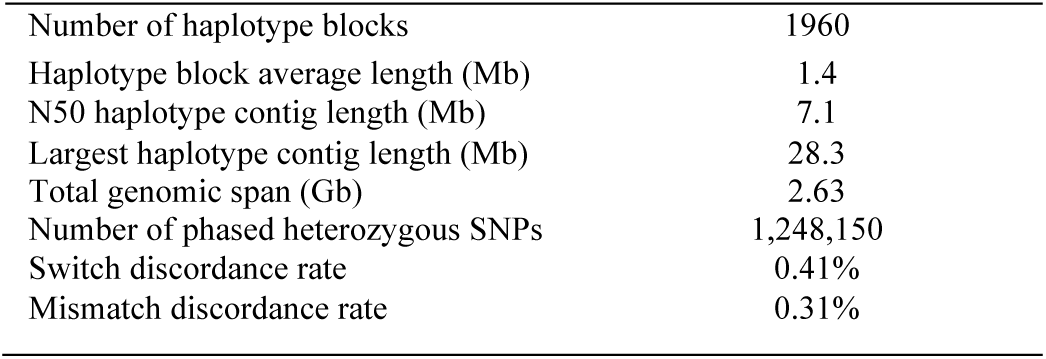
Summary of haplotyping performance.

|  |  |
| --- | --- |
| Number of haplotype blocks | 1960 |
| Haplotype block average length (Mb) | 1.4 |
| N50 haplotype contig length (Mb) | 7.1 |
| Largest haplotype contig length (Mb) | 28.3 |
| Total genomic span (Gb) | 2.63 |
| Number of phased heterozygous SNPs | 1,248,150 |
| Switch discordance rate | 0.41% |
| Mismatch discordance rate | 0.31% |

#### Error Rate of MDA, Library Preparation and Sequencing

To quantify the error rate, each SISSOR fragment was assigned to a haplotype by matching 2 or more heterozygous SNPs with over 80% accuracy in the assembled haplotype blocks (Fig. 2 and SI Appendix, Fig. S2E) and the number of mismatched bases in each pair of the haplotype-matched fragments was counted. The rate of mismatched bases between phased fragments with the same haplotypes obtained from between any cells is about 1.7 × 10^−5^ (SI Appendix, Table S5). The errors are very likely introduced by MDA, PCR-based library construction, and sequencing. The error rate is found to be consistent with that of other MDA-based sequencing methods (3, 14). The unphased fragments have fewer heterozygous SNPs due to their limited length, low coverage or being located within regions with low sequence complexity. The unphased fragments and reads outside the fragment boundaries were removed from downstream analysis.

Single-cell MDA is susceptible to trace amounts of DNA contamination in reagents and reaction vessels. The processing and amplification performed in the close environment of the microfluidic device with minimal flow paths and small volume of the microreactors minimizes contamination. This is reflected in the high mapping rate of sequencing reads and low mismatch rate we obtained (SI Appendix Table S2 and S5). Mutagenic damages associated with DNA oxidation could result in mutations which is usually prevalent in G to T mutations (15). We minimized oxidative DNA damage by including DTT (dithiothreitol) as a reducing agent in the denaturation solution. For the 3696 mismatches between the two complementary strands with a total ∼179 Mb (SI Appendix, Fig. S5), the rate of G to T mutation is only 2% while the ratio of transition to transversion mutations is 3.45. This indicates that oxidative damage was minimal or negligible in the SIS-SOR device.

#### Error Rate of Base Consensus in Phased Single-stranded DNA fragments

SISSOR technology can improve single-cell variant calling by matching same-haplotype strands, since variant calls that match between two complementary strands of DNA are of especially high confidence. However, it is not feasible to directly measure the accuracy of this approach for a single cell, since there is no reference genome for an individual cell with its *de novo* mutations. Instead, we provide an indirect estimate of the maximum possible error rate of same-haplotype strand matching in SISSOR technology, by noting that same-haplotype matching variant calls between strands in different cells do not carry cell-specific mutations (Fig. 3). We sampled haplotype-matched allele calls at the same genomic position using sequencing data in two chambers from different cells (cross-cell) and compared the matching consensus to the PGP1 reference. MDA errors, single cell *de novo* variants and haplotyping errors were removed by only considering the position with identical calls in two distinct cells (Fig. 3B, position 1-2). We expect that any unvalidated calls are either true errors, or true *de novo* SNVs found in the shared lineage of the individual cells.

**Fig. 3.**
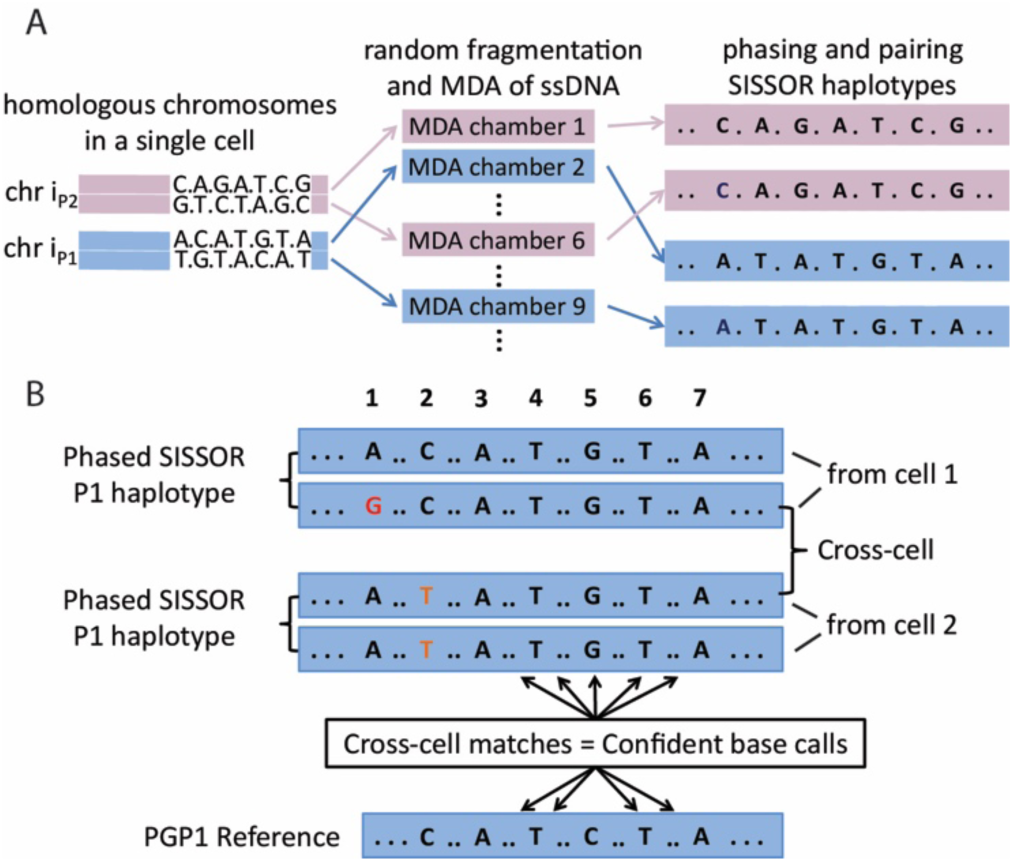
Error rate analysis of base consensus in phased SISSOR fragments. (A) Base sequences in single-stranded DNA fragments were constructed by variant calling of the mapped MDA products in each individual chamber and the complementary strands were identified by comparing the haplo-types of the single-stranded fragments from different chambers. (B) Matching variant calls in the contigs from the same haplotype between two cells (cross-cell), representing the PGP1-specific sequence, were validated by the PGP1/WGS reference. Common MDA and library preparation error was defined by the mismatches of variant calls between two matching phased haplotypes within the same cell (position 1). Single-cell *de novo* mutation was defined by matching variant calls between two matching phased haplotypes, together with matching variant call from at least one chamber in the other cell to the PGP1 reference (position 2). The rates of single chamber MDA-based sequencing error (10^−5^) and single-cell *de novo* mutation (10^−7^) were calculated for SISSOR. Cross-cell consensus, where *de novo* variants were removed, was defined by the matching variant calls between phased haplotypes in two different cells (position 3-7). The mismatch consensus to the PGP1 reference call (position 5) represented the discordance rate for SISSOR technology (10^−8^).

Phased haplotype fragments were randomly paired once between two cells at each unique haploid position (SI Appendix, Fig. S2F). 351 Mb unique cross-cells positions co-occurred with the consensus of Complete Genomics (CGI) (16) and Illumina WGS (17) reference of PGP1 (SI Appendix, Table S6). Only 19 matching cross-cells calls in SISSOR were discordant to the PGP1 reference (Fig. 4). Of these calls, 10 variants were found in the BAC reference and validated as true variants. Another 5 variants appeared in a third SISSOR chamber independent of the two haplotype-paired strands, indicating that these cases were not double errors in the phased strand consensus (Fig. 3B, position 5). Multiple appearance of the same variant confirmed that either this variant was previously undetected or only existed as *de novo* mutation shared by this cell lineage. Four variants remained unaccounted for. There is not sufficient information to determine whether that the variants are genuine *de novo* SNVs or are due to errors introduced by the SISSOR procedures. This bounds the overall sequencing error rate of the SISSOR technology below 1×10^−8^ (4 possible errors in 351 Mb). Further investigation showed that 1 of the 4 variants was supported by sequence from another chamber with a read depth of 4, which was lower than the threshold depth of 5. Another allele had PGP1 reference calls in the opposite haplotypes from the other chambers, which suggested the correct mapping and haplotyping at this location and further strengthened the possibility of these variants as true *de novo* calls.

**Fig. 4.**
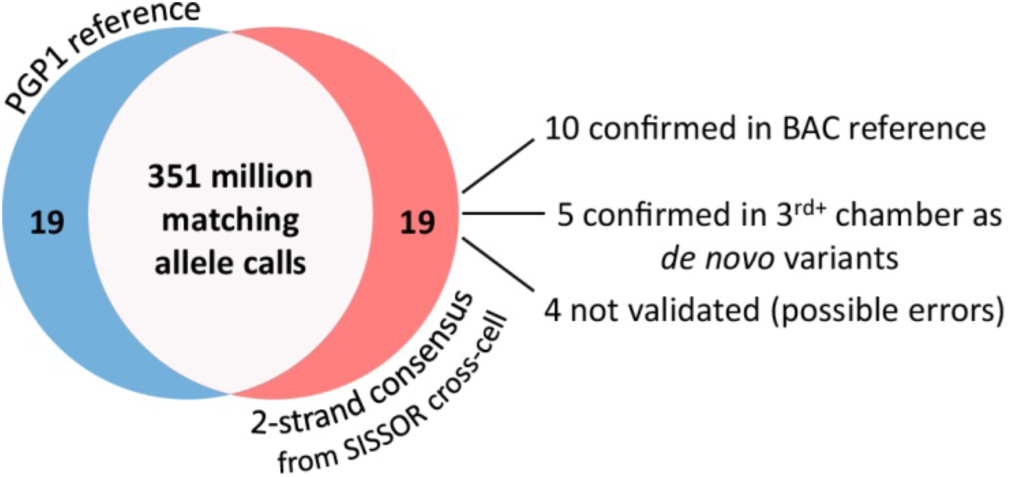
Differences between allele calls in PGP1 reference and SISSOR consensus. The consensus of CGI and Illumina WGS was used as the PGP1 reference. Positions lacking coverage in both PGP1 reference and SISSOR consensus were discarded.

#### Direct Measurement of *de novo* Variants in a Single Cell

We expect sequencing accuracy equivalent to the strand-strand consensus in the same cell, assuming error rate was identical in all SISSOR libraries. Number of matched SNVs within a single cell was presumably higher than cross-cell matches because of the true *de novo* variants in a single cell. In comparison to bulk sequencing, individual mutations from each single cell are masked and undetected by the consensus of many other cells. In contrast, the consensus of phased SISSOR fragments from each single cell detected a total of 68 possible *de novo* SNVs (SI Appendix, Table S7). None of these were found in the other PGP1 libraries prepared using induced pluripotent stem cell and fibroblast (16), where their combined consensus was about 0.1% different than lymphocyte only. Correct PGP1 variant calls were found in 28 (∼41%) SNVs, such as position 2 of cell 1 in Fig. 3B where novel calls were discovered only in cell 2. 7 (∼10%) base-calls were found to have the correct PGP1 reference base called in the opposite haplotype within the same cell. 17 (∼25%) were found to have the correct PGP1 reference base called in both the opposite haplotype and the other cell. Many of these variants appear to be real since correct PGP1 reference base calls suggest correct mapping at the identical positions. The remaining 16 SNVs lacking coverage from the other cells are potential false-positive errors. However, all but two unsupported SNVs were presumably undetected *de novo* variants because of the lower error in SISSOR technology.

#### Direct Resolution of the HLA Haplotypes

Haplotype of the HLA genes has been determined using SISSOR. The highly polymorphic HLA loci are distributed within a ∼5 Mb region on chromosome 6 (28.5M-33.5M) and are important to the immune system. We obtained a single SISSOR haplotype block of ∼18.2Mb spanning the entire HLA region. This allows for the phasing of the four classical HLA genes. We selected the sorted BAM files from individual SISSOR fragments that correspond to the HLA haplotype and further determined the best genotypes of HLA genes via the tool bwakit (under github “lh3/bwa”). Top matches of all four HLA genes were identified in one of the two HLA haplotypes: HLA-A*02:01:01:01, HLA-B*51:01:01, HLA-C*05:01:01:01, HLA-DRB1*13:01:01 (SI Appendix, Fig. S6).

## Conclusion

In this study, we have demonstrated whole-genome sequencing of single cells using the SISSOR technology in which megabase-size single-stranded DNA fragments from the homologous chromosome pairs of a single cell are partitioned into multiple compartments for independent amplification and sequencing. With two-strand consensus and long sub-haploid fragment assembly, our approach can simultaneously provide higher per base sequencing accuracy and longer haplotypes than other techniques where similar single cell amplification and sequencing platforms are employed. Unlike the long fragment read method (18), in which errors are reduced by comparing consensus calls from multiple single-stranded libraries from many cells, our SISSOR technology makes use of the consensus of the two complementary strands from only a single cell for error corrections. The long SISSOR fragments also facilitated long-range haplotype assembly with short-read sequencing data acquired from isolated single cells, without the needs for extensive cloning (8) and multiplying cells in culture (10). This makes possible high sequencing accuracy using rare single cells or non-dividing cells from primary tissues, such as adult neurons (19). Using a reference genome, the accuracy of two-strand variant consensus calls was confirmed to be better than 1 error in 100 million bases, which is a dramatic improvement from 2000 errors in 100 million bases obtained using double-stranded sequencing library. The accuracy exceeds what is required for detecting the potential genetic variations between single cells (10^−7^). Our current implementation of the SISSOR technology still has limitations, including the lack of integrated sequencing library preparation and scalability for parallel processing and sequencing of multiple single cells. These limitations can be addressed by designing microfluidic processor with polymer barriers to enable single-chamber multistep processing required for on-chip preparation of encoded sequencing libraries (20), and integrating the water-in-oil droplet-based approach to enable parallel genome sequencing of multiple single cells. We anticipated diverse research and clinical needs will be enabled by highly accurate genome sequencing and haplotyping of single cells.

## Materials and Methods

### Fabrication of microfluidic processors

The PDMS microfluidic devices with domed flow channels at the top layer and 25 μm thick valve control layer at the bottom were fabricated using soft lithography. Both layers are bonded together and then to a glass slide using standard PDMS techniques (20, 21). The mold for the fluidic channels with multiple heights was fabricated by repeating the soft lithography process. All domed channels (12 μm and 20 μm) were fabricated using photoresist AZ 12XT-20PL-10 (AZ) followed by reflowing to produce the domed structure. The 60 μm tall rectangle MDA chambers are fabricated using SU8 2050 (MicroChem). The molds were passivated with tride-cafluoro-1,1,2,2-tetra-hydrooctyl-1-1trichlorosilane (Pfaltz and Bauer, Inc.) prior to casting PDMS. The PDMS layers were fabricated using Sylgard 184 (Dow Corning). A 5:1 mixture (part A: part B = 5 :1) was poured onto the mold for the top fluidic layer. A 20:1 mixture (part A: part B = 20:1) was spin-coated onto the mold for the valve control layer at 1500 rpm for 45 seconds. After both PDMS layers were cured on the molds for 25 min at 65°C in an oven, the fluidic layer was peeled off and access holes were created using a 0.75 mm diameter biopsy punch (Ted Pella, Inc.). The fluid layer was then aligned and laid onto the thin valve control layer. The two PDMS layers were bonded for 4 hours at 65°C in an oven. The bonded layers were peeled off and the valve connecting holes were punched. The surface of the valve layer and a cover glass (75 mm × 50 mm × 1.0 mm) were treated with oxygen plasma in a UV-ozone cleaner (Jelight Company, Inc.) for 4 min, and then bonded together for 10 hours at 65°C in an oven. A photograph of a working device is shown in Fig. S1 in SI Appendix,.

### Partitioning, amplification and sequencing of single-cell genome

On chip amplification was performed with a modified protocol of the Nextera Phi29 kit. After filling all valve lines with pure filtered water (18 M·cm), all MDA chambers were incubated for 15 minutes with 0.1% BSA, 35 mM random hexamers and 16 μM dNTPs, then purged and dried with air flow. The entire device was then sterilized for 15 minutes in an ultraviolet crosslinker (Longwave UV Crosslinker, UVP). A single cell suspended in PBS buffer was immediately loaded in the capture chamber, then flushed and filled with alkaline lysis solution (ALS) (400 mM KOH, 10 mM DTT and 1% Tween20) in the mixing chamber. Cell lysate was mixed with the lysis solution in the ring mixer for 10 minute using the PDMS peristaltic pump operating at 5 Hz, and then loaded in the 24 partition chambers by pushing with air. A neutralization solution (NS) (400 mM HCl, 600 mM Tris-HCl and 1% Tween20) and Nextera MDA mastermix (1x MDA buffer, 84 μM 3’ phosphorylated random hexamers, 2 mM dNTPs, 150 μM dUTP and 0.84 μg of Phi29 DNA polymerase) were loaded into a channel parallel to the partition chambers, and all partitioning valves were closed along the NS line and partitioning chambers. The reaction mix was pushed into the individual MDA chamber and the device was incubated at 30°C for 15 hours to carry out the MDA reactions. All the feed lines were flushed with air and washed with TE buffer. After amplification, the MDA products in each chamber was pushed into a pipet tip with 1x TE buffer at the individual outlet and 5 μl was collected. The samples were incubated at 65 °C for 15 minutes to inactivate Phi29 DNA polymerase. Construction of barcoded sequencing libraries and Illumina sequencing were performed as described by Peters et al. (18) and Adey et al (22) and further described in the supplementary methods.

### Mapping sequencing reads

Human genome extension hs37d5 (GRCh37+decoy) from the 1000 Genomes Project was used as the reference for mapping. All reads were mapped with BWA-MEM (23). Paired-end 100 bp reads were mapped as two single-end reads. All reads in the fastq format were aligned to the human genome reference in the fasta format. Alignments were obtained in the SAM/BAM format (24). For BWA-MEM alignment, the “mem#8211;M” command was applied to BWA (version 0.7.10).

## Acknowledgements

This work was supported by NIH grant R01HG007836 to (KZ & XH), and was performed in part at the San Diego Nanotechnology Infrastructure (SDNI) of UCSD, a member of the National Nanotechnology Coordinated Infrastructure, which is supported by the National Science Foundation (Grant ECCS-1542148)

### Conflict of interest statement

XH and KZ are listed as inventors for a patent application related to the method disclosed in this manuscript. KZ is a co-founder and equity holder of Singlera Genomics Inc. VB is a cofounder, has an equity interest, and receives income from Digital Proteomics, LLC. The terms of this arrangement have been reviewed and approved by the University of California, San Diego in accordance with its conflict of interest policies.

